# An open-source and wearable system for measuring 3D human motion in real-time

**DOI:** 10.1101/2021.03.24.436725

**Authors:** Patrick Slade, Ayman Habib, Jennifer L. Hicks, Scott L. Delp

## Abstract

Analyzing human motion is essential for diagnosing movement disorders and guiding rehabilitation interventions for conditions such as osteoarthritis, stroke, and Parkinson’s disease. Optical motion capture systems are the current standard for estimating kinematics but require expensive equipment located in a predefined space. While wearable sensor systems can estimate kinematics in any environment, existing systems are generally less accurate than optical motion capture. Further, many wearable sensor systems require a computer in close proximity and rely on proprietary software, making it difficult for researchers to reproduce experimental findings. Here, we present OpenSenseRT, an open-source and wearable system that estimates upper and lower extremity kinematics in real time by using inertial measurement units and a portable microcontroller. We compared the OpenSenseRT system to optical motion capture and found an average RMSE of 4.4 degrees across 5 lower-limb joint angles during three minutes of walking (*n* = 5) and an average RMSE of 5.6 degrees across 8 upper extremity joint angles during a Fugl-Meyer task (*n* = 5). The open-source software and hardware are scalable, tracking between 1 and 14 body segments, with one sensor per segment. Kinematics are estimated in real-time using a musculoskeletal model and inverse kinematics solver. The computation frequency, depends on the number of tracked segments, but is sufficient for real-time measurement for many tasks of interest; for example, the system can track up to 7 segments at 30 Hz in real-time. The system uses off-the-shelf parts costing approximately $100 USD plus $20 for each tracked segment. The OpenSenseRT system is accurate, low-cost, and simple to replicate, enabling movement analysis in labs, clinics, homes, and free-living settings.

## I. Introduction

Researchers and clinicians measure kinematics during human movement to attain an understanding of diseases and rehabilitation, to improve the design of assistive devices, or to uncover the mechanisms by which athletes achieve peak performance. Estimating kinematics in real-time is important for many real-world applications of movement analysis, such as rapid biomechanics assessment for injury prevention, corrective feedback for at-home rehabilitation, or real-time control of assistive devices to improve mobility.

Motion capture systems use sensors to objectively measure the motion of body segments and estimate joint angles and other kinematic quantities. Motion capture systems typically rely on either stationary sensors such as cameras placed throughout a room or wearable sensors such as inertial measurement units (IMUs) attached to the body. Systems that measure joint kinematics with less than 2 degrees root-mean-square error (RMSE) are considered to be excellent and less than 5 degrees RMSE to be acceptable for many applications [1], although accuracy requirements will depend on the application.

Optical motion capture is the current industry standard. Careful calibration can enable these camera systems to track the position of markers with millimeter accuracy. However, capturing human motion adds additional sources of error such as skin motion artifacts and occluded markers [2], which can increase the average RMSEs for joint angles by 1 to 3 degrees [3]. The use of optical motion capture is limited by the high cost of the equipment and the requirement of a fixed capture volume.

Wearable systems use portable sensors, most commonly IMUs, worn on the body to capture motion in many environments. Modern IMUs measure linear acceleration, angular velocity, and the local magnetic field and are compact, lightweight, and low-cost. Joint kinematics can be computed with a variety of methods including integration [4], constrained orientation [5], statistical models [6], or musculoskeletal simulation [7]. Commercial IMU motion capture systems can estimate joint kinematics with less than 5 degrees RMSE compared to optical motion capture during brief bouts of activities including walking, jumping, and squatting [8]. Most current wearable systems estimate kinematics offline due to the significant number of processing steps and computation requirements. Computing kinematics in real-time is necessary for providing rapid results or diagnoses as well as applications that require feedback to the wearer, such as correcting an exercise or improving an athletic maneuver.

Beyond computing kinematics in real-time, wearable systems should meet several additional criteria to be effective across a wide range of real-world scenarios. First, the accuracy and frequency of estimation must be sufficient for the application of interest. Systems that require off-board computation, such as a laptop or desktop, to provide real-time estimates restrict the motion capture volume to the range where the sensors can communicate with the computer. Thus, on-body computation is preferred for use in a wide range of environments. A wearable motion capture system for real-world applications also requires robustness to environmental changes such as magnetic field and temperature. Design guidelines for wearable medical devices suggest devices should be lightweight, low-cost, and easy to don and doff for extended use [9]. In addition, systems with open-source software, simple assembly procedures, and a straightforward calibration method would reduce barriers to use.

Existing real-time wearable systems span a range of features (Table I). All of these systems require off-board computation, which increases the overall system cost and limits motion capture to the range in which sensor data can be passed to the computer. Further, existing systems either use custom hardware and are not open-source, use commercial hardware and are expensive, or compute kinematics for a fixed set of joints for a specific application. While current commercial systems meet some of the desired specifications for real-time wearable systems such as accurately estimating joint kinematics, high computation frequency, and flexibility in the number of tracked body segments, they are expensive and require the user to stay within approximately 40 feet of a desktop or laptop computer.

**TABLE I.**
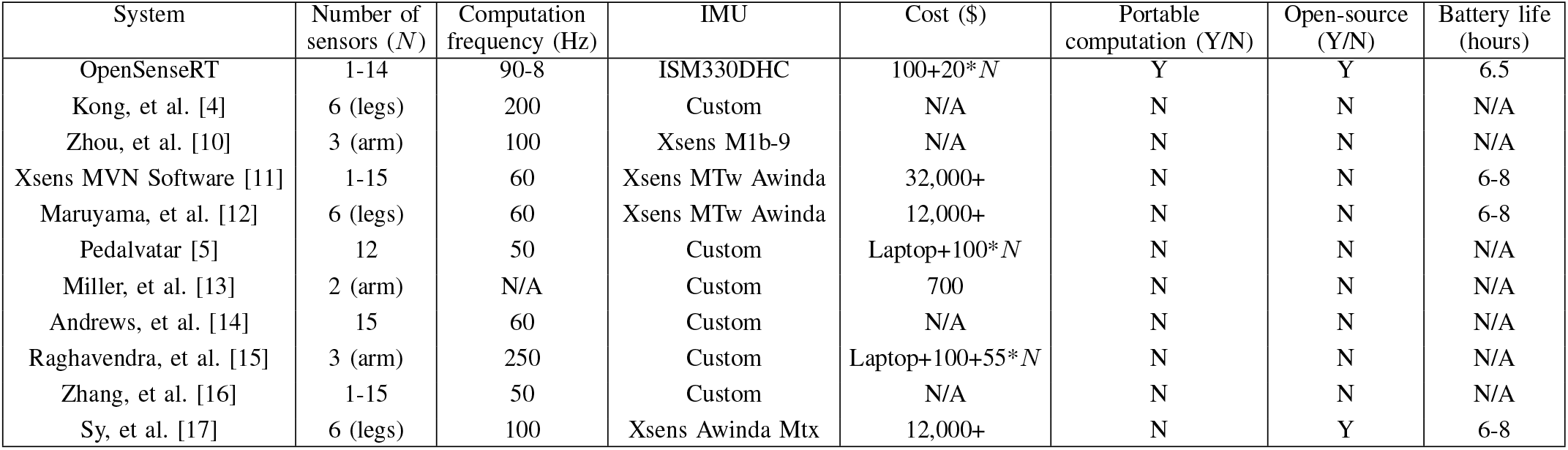
Comparison of wearable motion capture systems that compute real-time kinematics

To help overcome the current barriers to widespread use of real-time wearable systems, we developed the OpenSenseRT system, a low-cost, open-source, and fully wearable system for computing real-time kinematics. We assessed accuracy by comparing the OpenSenseRT system to optical motion capture for a range of upper and lower extremity movements, seeking to achieve an RMSE of 5 degrees or less for 3D joint angles. To make the system useful in a wide range of applications we used off-the-shelf components and enabled customization to estimate any combination of lower and upper extremity motions (assuming one sensor worn on each tracked segment). We have provided the software, a list of components, instructions for assembly and use, and examples of activities measured with the OpenSenseRT system [18].

## II. Methods

### A. Open-source system hardware

The system hardware is built from components that are inexpensive and can be assembled without tools or soldering. The components include IMUs, a microcontroller, Qwiic connector cables, a button (to start and stop recording), and a rechargeable battery (Fig. 1). At the time of publication, parts to build the system cost approximately $100 plus $20 for each tracked body segment. The assembly procedure consists of assembling a case for the microcontroller and fan, attaching this case to a belt to be worn on the pelvis, and connecting the sensors to the microcontroller. A complete bill of materials, suggested vendors, written assembly instructions, and video of the assembly procedure are provided in a user’s guide to simplify replication of the system [18].

**Fig. 1.**
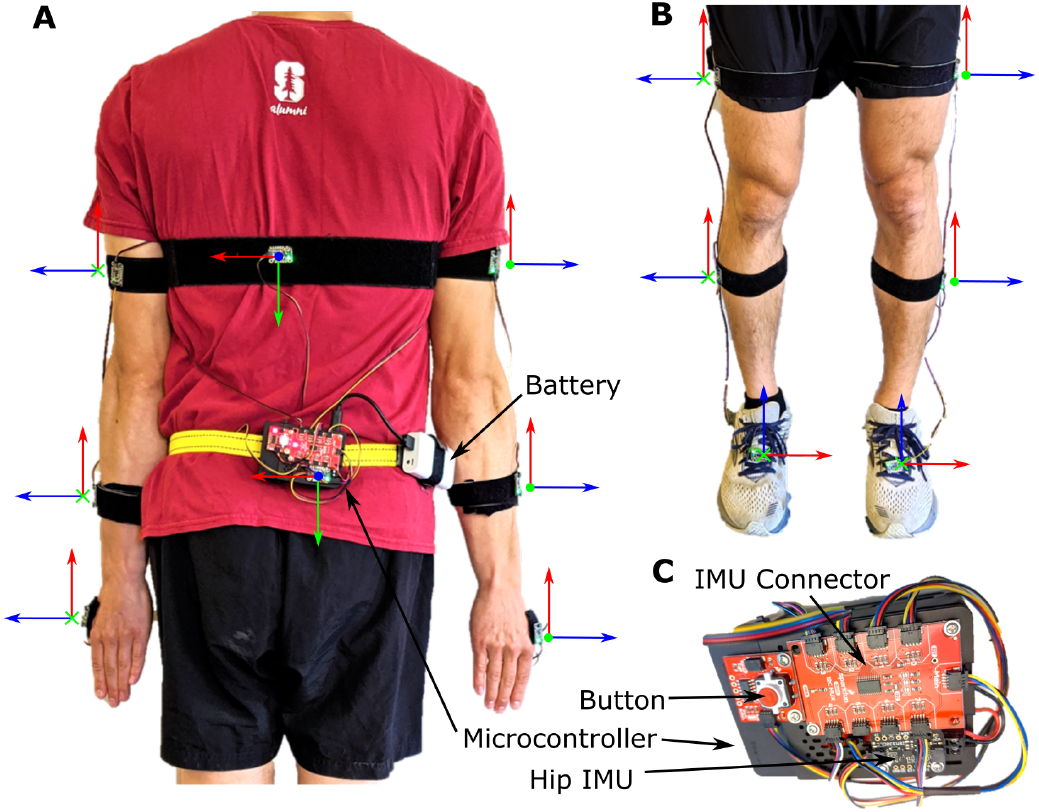
The components and default IMU orientations of the OpenSenseRT System. (A) An IMU on the pelvis is required and acts as the base in order to compute the relative orientation of other sensors. The OpenSenseRT System accommodates a variable number of additional IMUs to customize which kinematics are measured. To monitor movement of the upper body, three IMUs may be placed on each arm (on the upper arm, forearm, and hand). An additional IMU can be placed on the torso. The orientation frame with axes shown in red, green, and blue is used to orient the ***x, y***, and ***z*** axes defined on each IMU. These individual body frames should align with the world reference frames of the fore-aft, mediolateral, and vertical axes, while the subject’s joint segments are aligned in a neutral standing (or other known) position. (B) The lower-limb IMU placements also require the pelvis IMU as a base and can include up to three IMUs on the thigh, shank, and foot of each leg. OpenSenseRT allows for 1 to 14 of these IMUs to be sampled in a custom configuration. (C) A zoomed in view of the system components shows the microcontroller, battery, button (for starting and stopping recordings), IMU connector, and pelvis IMU.

The microcontroller and IMUs were selected because they meet computational requirements for real-time estimation of joint kinematics. The microcontroller is a Raspberry Pi 4b+ with 4 gigabytes of RAM and a quad-core microprocessor (Raspberry Pi Foundation). This amount of RAM is required to build the OpenSim software [19] that performs the inverse kinematics. The quad-core microprocessor allows for multiple threads to run in parallel, necessary for the system’s software architecture discussed in the next section. The IMUs are ISM330DHCX breakout boards (Adafruit) which contain an accelerometer and gyroscope. These IMUs measure accelerations up to 16 times gravity and angular velocities up to 4000 degrees per second, sufficient to capture dynamic activities like running. The IMU sampling rate is variable with a maximum rate of 6.7 kHz. The gyroscope has low measurement noise with a standard deviation of 0.1 degrees per second, which helps to minimize drift in the measurements. The IMU breakout board includes connectors to simplify wiring the IMUs to the microcontroller with pre-made cables. We recommend gluing the pre-made cables into connectors on the IMU and microcontroller to improve the strength of the connections.

The hardware system specifications are designed to estimate joint kinematics for extended periods in free-living settings. The total weight of the system is 408 grams, with approximately half due to the rechargeable battery. The rechargeable battery allows continuous recording for approximately 6.5 hours per charge and can be recharged in about 2 hours or replaced for continual use. The SD card in the microcontroller is able to store approximately 100 hours of recordings at a 30 Hz computation rate. A button and blinking LED provide control and feedback to the user to start and stop recordings.

### B. Open-source software architecture

The open-source software requires user inputs to define which body segments are tracked with IMUs and uses a two phase procedure to estimate kinematics: initial calibration and iterative computation of joint kinematics (Fig. 2). The microcontroller’s SD card contains the operating system for the OpenSenseRT system. The user-defined parameters are input to the system by editing a text file on the SD card. Step-by-step instructions to edit this text file and additional instructions to customize the software are provided in the user’s guide [18].

**Fig. 2.**
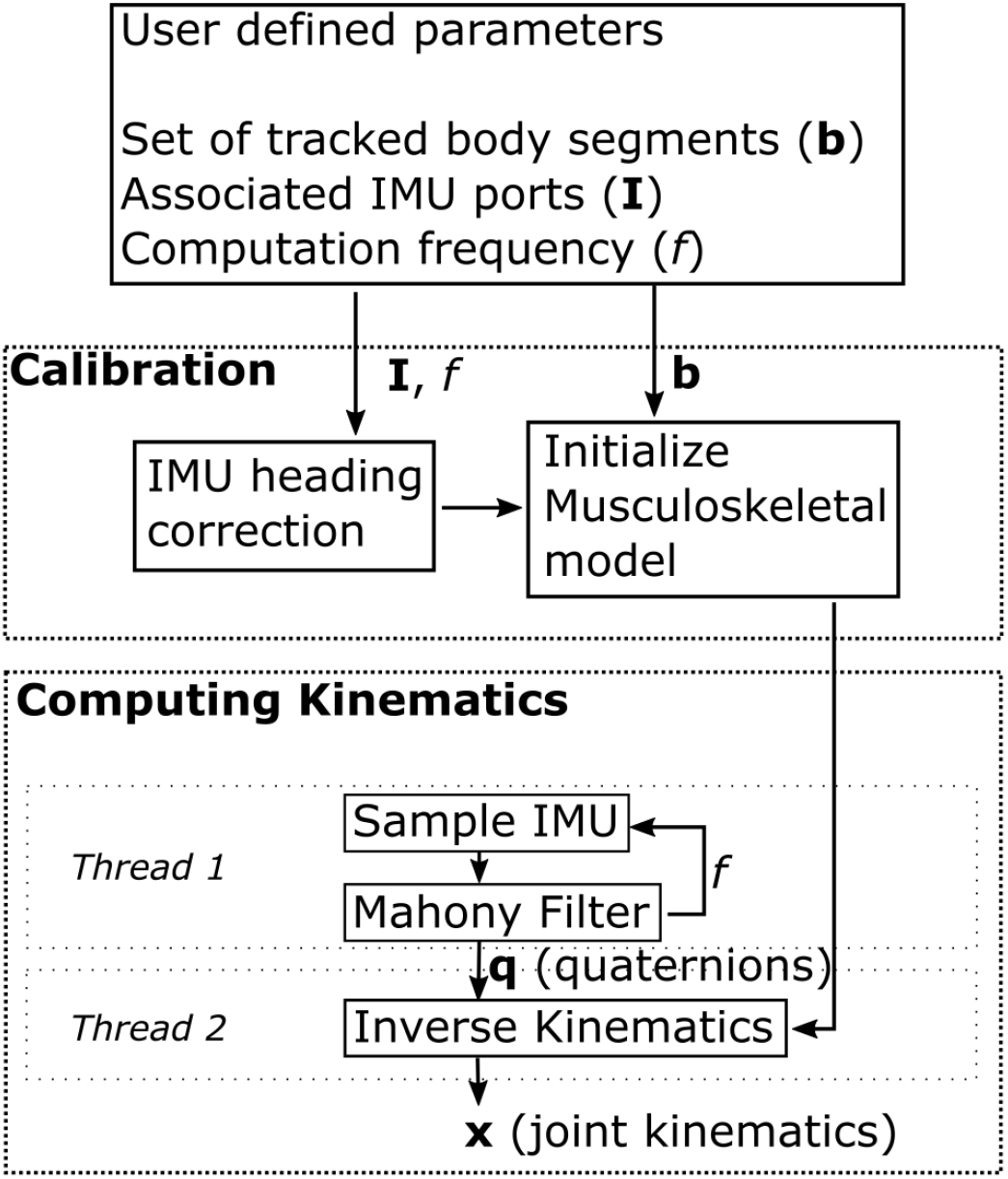
Flowchart for computing joint kinematics. The user defines the set of body segments to track, the IMU ports associated with the segments, and the computation frequency. The calibration process consists of a heading correction for the given IMUs and initializing the pose of the musculoskeletal model. The calibrated model and initial pose are passed to the inverse kinematics solver. One thread on the microcontroller records raw IMU data and computes orientations for each body segment using a Mahony Filter at each step. A second thread takes these orientations and the musculoskeletal model to solve the inverse kinematics, estimating joint kinematics at each time step.

The software requires a calibration of the gyroscopes in the IMUs during their first use. Gyroscope measurements are taken for 10 seconds and averaged to remove any offset, which prevents drift due to sensor bias. These gyroscope bias values are stored and recalled for any future recordings with the same set of sensors.

Before each recording session, the system must be calibrated; we assume a standing pose and a set of known IMU orientations with respect to each body segment. The subject stands in a neutral position with their arms at their sides, with all IMUs aligned as shown in Fig. 1. Then the heading angle of the IMUs with respect to the anterior-posterior axis of the musculoskeletal model is averaged across all IMUs. The individual heading of each IMU is corrected by adding an offset to match this average heading angle. This heading correction removes small deviations in the orientations between the IMUs during the subject’s standing pose and the initial generalized coordinates of the musculoskeletal model used to estimate kinematics.

Once calibrated, the system uses the inverse kinematics solver of OpenSim to compute new joint kinematics at each timestep based on the IMU orientations. An IMU on the pelvis is required and acts as the base to compute the relative orientation of other sensors. The microcontroller uses two separate threads to compute joint kinematics at each step. The first thread samples each IMU and converts the raw accelerometer and gyroscope readings into a quaternion containing the orientation by using a Mahony Filter with a proportional gain of 1 and integral gain of 0.3 [20]. The proportional gain was left to the recommended default value of 1 and an integral gain of 0.3 was chosen empirically to allow the initial IMU orientations to reach a steady-state value within 3 seconds. Magnetometer measurements were not used to compute the quaternions because they are sensitive to changes; for example, when near electric motors or large metal objects.

The second thread uses these orientations and the musculoskeletal model as inputs to the inverse kinematics solver included in OpenSim version 4.2 [19] to compute the generalized coordinates of the model (i.e., joint angles and joint angular velocities). The inverse kinematics solver uses the generalized coordinates of the musculoskeletal model at the previous time step as an initial guess. The new set of generalized coordinates are computed by minimizing the error relative to the IMU orientations from the current step, converting to an axis-angle representation, and then minimizing the difference in the joint angles. The axis-angle representation defines orientation in three dimensions with a unit vector, or axis, and a single angle about this axis. These steps are repeated to estimate the kinematics as measurement moves forward in time. Using two threads reduces the delay in computing kinematics, allowing for higher computation frequencies. The kinematics are computed from the relative orientations, eliminating complex model scaling steps required to accurately process motion capture data that relies on the absolute positions of markers placed on the body.

Estimating kinematics in real-time required several modifications to the musculoskeletal model and inverse kinematics settings. We simplified a previously published full-body musculoskeletal model [21] by removing muscles and locking unused joints, including the subtalar and metatarsophalangeal joints. The inverse kinematics solver used a weight on the joint constraints of 10 and a solver tolerance of 0.001. These values were selected through trial and error in pilot testing of the device to achieve both accurate and fast solutions.

The OpenSenseRT system can estimate kinematics online or offline. The online mode uses one or more of the microcontroller cores to compute kinematics. The offline mode only samples the IMUs, performing the inverse kinematics computation later. The offline mode provides much faster sampling frequencies by removing the need to compute kinematics in real-time. Higher frequency estimates of joint kinematics improve the accuracy when estimating dynamic motions such as running and may be preferred for applications without real-time constraints.

### C. Experimental comparison to optical motion capture

We compared estimates of joint kinematics from the OpenSenseRT system to estimates based on optical motion capture (Optitrack) for a range of conditions including: walking, running, a Fugl-Meyer upper-limb task [22], and a task measuring the range of motion of the trunk. The motion capture was recorded at 100 Hz. Lower-limb experiments used sixteen markers, including markers bilaterally on the anterior and posterior superior iliac spines, as well as markers on the right leg at the 2nd and 5th metatarsal heads, calcanei, malleoli, and femoral epicondyles, a four-marker plate on the shank, and a three-marker plate on the thigh. Upper-limb experiments used 14 markers bilaterally on the anterior and posterior superior iliac spines, as well as markers on the right arm at the C7 vertebrae, right clavicle, anterior and posterior shoulder, lateral shoulder, lateral and medial epicondyles of the humerus, radial and ulnar styloid processes, and the body of the 5th metacarpal. Healthy young adults (*n* = 5, 3 men and 2 women; age = 25.0 ± 1.8 yr; body mass = 63.0 ± 10.3 kg; height = 1.70 ± 0.07 m) were recruited. All subjects were volunteers and provided written informed consent before completing the protocol IRB-17282 approved by the Stanford University Institutional Review Board. Only one subject completed the running condition, but all subjects completed all other conditions. The walking condition lasted 3 minutes on a treadmill at 1.25 m/s. The running condition lasted 1 minute on a treadmill at 4.0 m/s. The Fugl-Meyer upper-limb and trunk range of motion conditions were completed ten times to create a cyclic upper-limb task for analyzing an average activity cycle in a similar manner to a gait cycle [23]. The Fugl-Meyer upper-limb task simulated cutting by having the subject pick up a knife from a table, perform a cutting motion, and place the knife back on the table [24]. The range of motion of the trunk task had subjects reach their comfortable range of motion for trunk flexion, then rotation, then bending [25].

The optical motion capture data was input to the OpenSim software [19] to estimate the ground truth kinematics. The optical motion capture data was processed following best practices to ensure proper model scaling, an accurate initial pose, and precise tracking of the marker positions using OpenSim’s inverse kinematics solver [26]. The joint kinematics computed with the OpenSenseRT system utilized the previously described open-source hardware and software.

While our validation focused on joint kinematics, the OpenSenseRT system can compute any quantity that is derived from the kinematic state of the model. For demonstration purposes, we computed foot progression angle. We first computed virtual marker positions on the heel and second metatarsal head, using OpenSim’s analyze tool offline. Then the foot progression angles were computed from these virtual marker positions with a previously defined approach [27]. The OpenSenseRT system can perform kinematic analyses in real time, but would require custom development for a given application.

## III. Results

### A. Accuracy of estimated joint kinematics

The OpenSenseRT system estimated lower-limb joint kinematics during walking with an average RMSE of 4.4 degrees compared to kinematics computed from optical motion capture data. Average joint kinematic curves from the OpenSenseRT system were similar to values from optical motion capture for both walking (Fig. 3A) and running (Fig. 3B). The joint kinematic curves from OpenSenseRT had a larger standard deviation than optical motion capture for the running trials, indicating the system was more susceptible to fluctuations during faster motion or higher impact activities. The absolute RMSE averaged across all joints drifted from 3.5 to 6 degrees after 3 minutes, although most angles stayed below a RMSE of 5 degrees for the first minute of data capture (Fig. 4). Hip rotation had the largest RMSE among the joint angles measured, but also has larger errors relative to other joints when measured with optical motion capture [3].

**Fig. 3.**
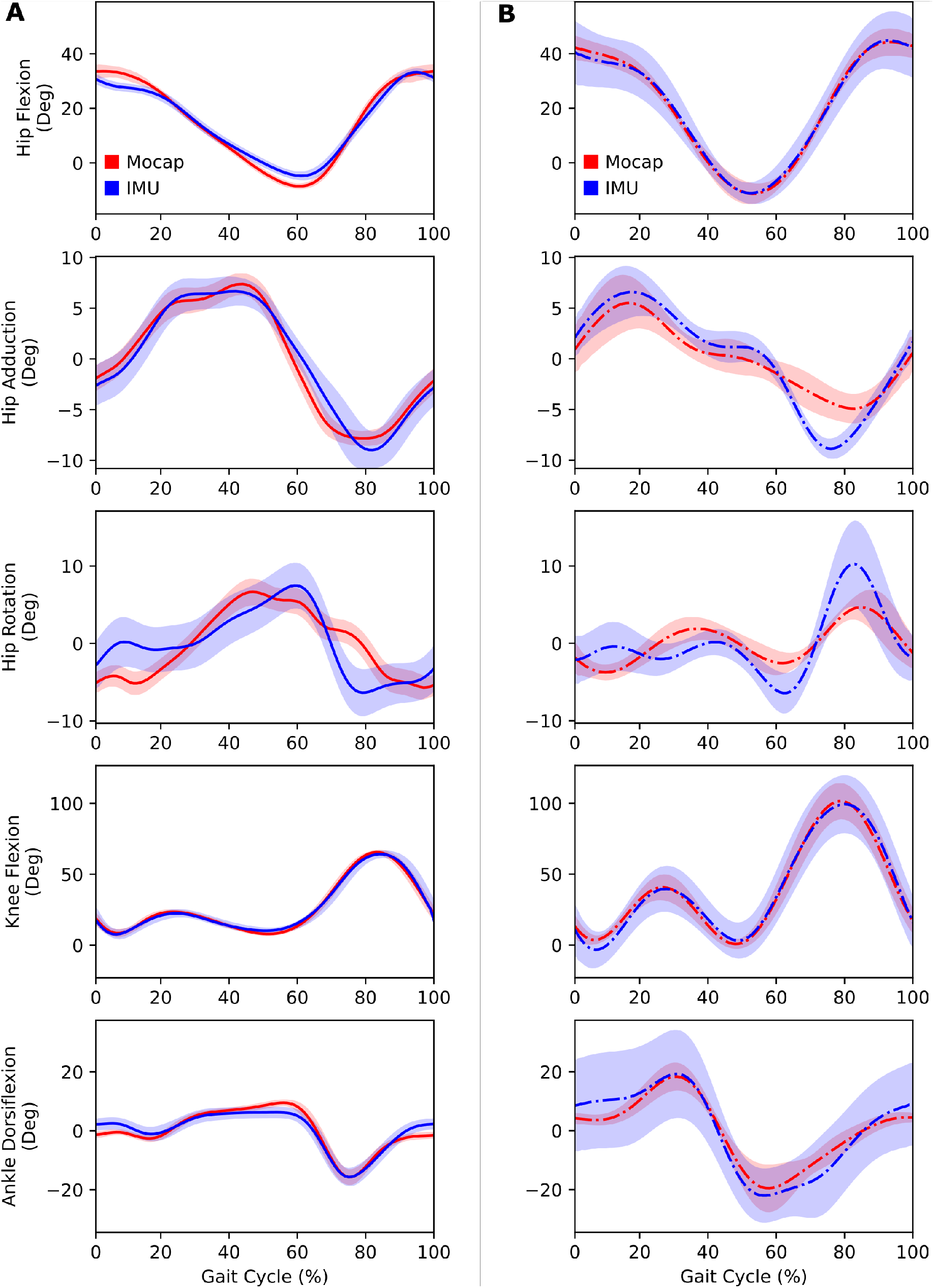
Lower-limb joint kinematics computed with the OpenSenseRT system compared to optical motion capture. (A) Joint kinematics during walking at 1.25 m/s were computed in real-time at 30 Hz with the IMU system for five subjects. The error bands represent one standard deviation of the RMSE across all subjects. (B) Joint kinematics during running at 4.0 m/s were computed offline (represented with the dash-dot line) at 100 Hz with the IMU system for a single subject. The error bands represent one standard deviation of the RMSE across all gait cycles for the single subject. Running requires a faster computation rate than the IMU system can perform consistently in real-time, although it can be analyzed in real-time over short bursts by using all cores.

**Fig. 4.**
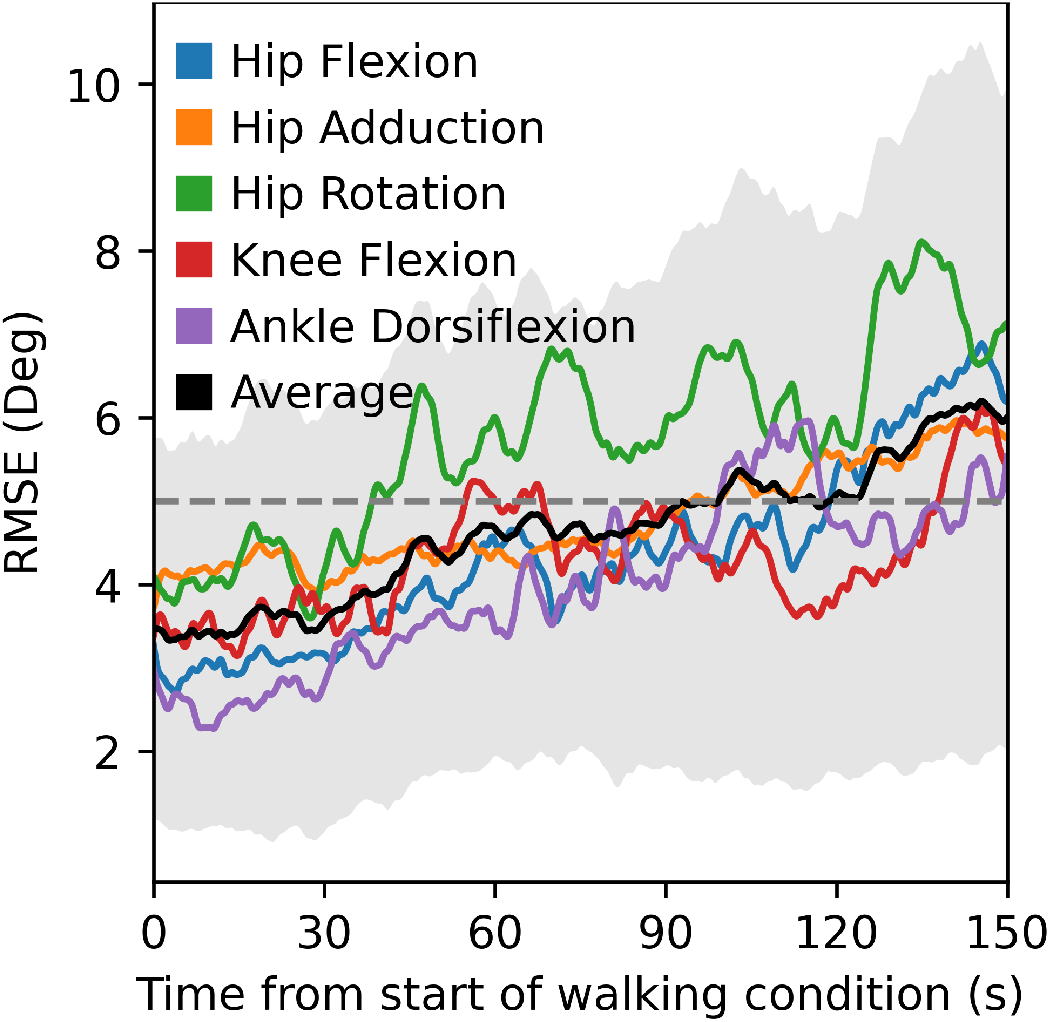
RMSEs for joint kinematics computed with the IMU system compared to optical motion capture over time for walking. The RMSE increased from approximately 3.5 to 6 degrees over the first 150 seconds of the condition. Hip rotation had the largest RMSE and drift. The black line and grey error band represent the average and standard deviation in RMSE across all subjects and joints.

The OpenSenseRT system estimated upper-limb joint kinematics with an average RMSE of 5.6 degrees compared to kinematics computed from optical motion capture. The individual upper-limb joint angles had average RMSEs between 2.5 and 10 degrees (Fig. 5). Average joint kinematic curves from the OpenSenseRT system were similar to values from optical motion capture for the upper-limb task (Fig. 6A). The joint kinematics during the upper-limb task had larger standard deviations than other conditions for both OpenSenseRT and optical motion capture, because the activity cycles occurred over a longer time period and the movement was more variable across repetitions and subjects. The trunk joint kinematics estimates during the trunk range of motion task show the joint kinematics curves from the OpenSenseRT system and optical motion capture have similar means and standard deviations (Fig. 6B).

**Fig. 5.**
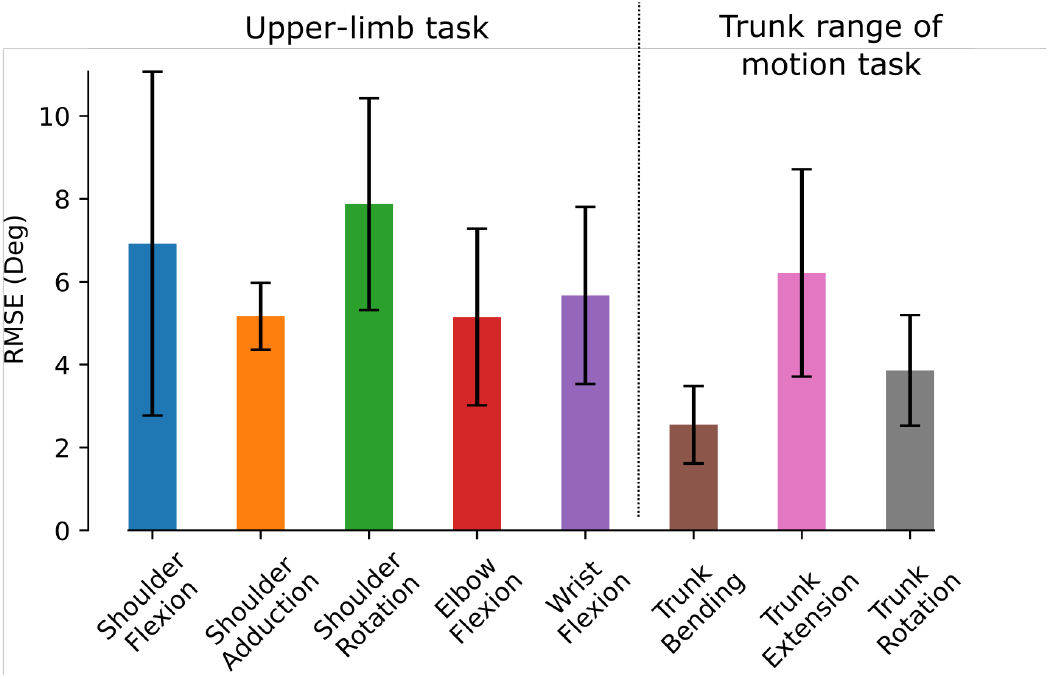
The average RMSE across each joint for the upper-limb task and the trunk range of motion task. The error bars represent one standard deviation of the RMSE across all subjects.

**Fig. 6.**
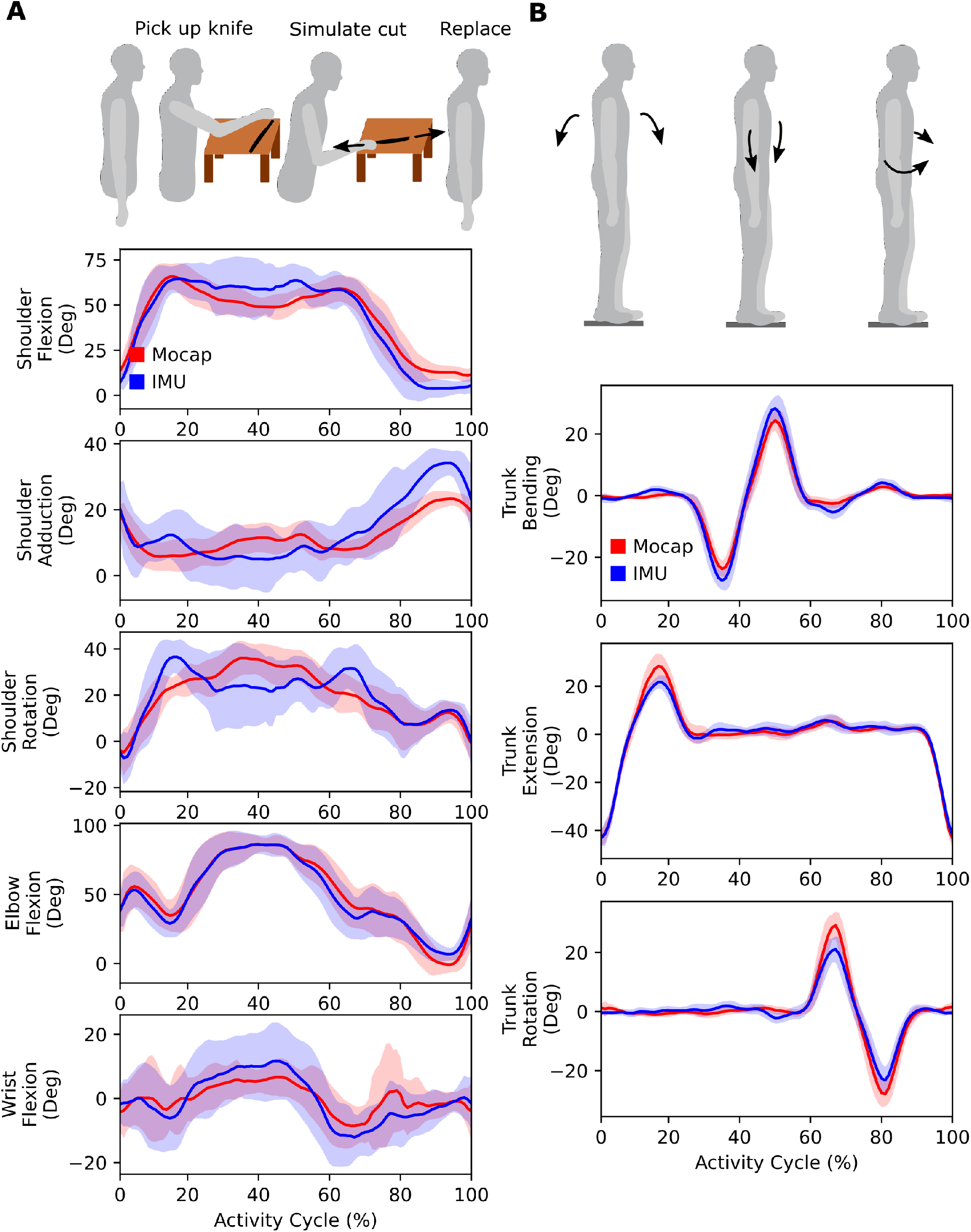
Upper-limb joint kinematics computed with the IMU system compared to optical motion capture. (A) An upper-limb task from the Fugl-Meyer assessment simulated cutting food. Subjects picked up a cutting utensil bent over a dish, and performed a cutting motion before placing the utensil back down. The task was repeated ten times to emulate an upper-limb “activity cycle” to look at average kinematics for one subject. (B) In the trunk range of motion task, subjects performed trunk flexion, bending, and then rotation. In both (A) and (B), the bands represent one standard deviation of the RMSE across all subjects.

Foot progression angle was computed to evaluate an example of a clinically relevant metric that is derived from the generalized coordinates of the model. The estimated foot progression angle for a single individual showed similar trends to the values computed based on optical motion capture data as the angle was varied during walking (Fig. 7).

**Fig. 7.**
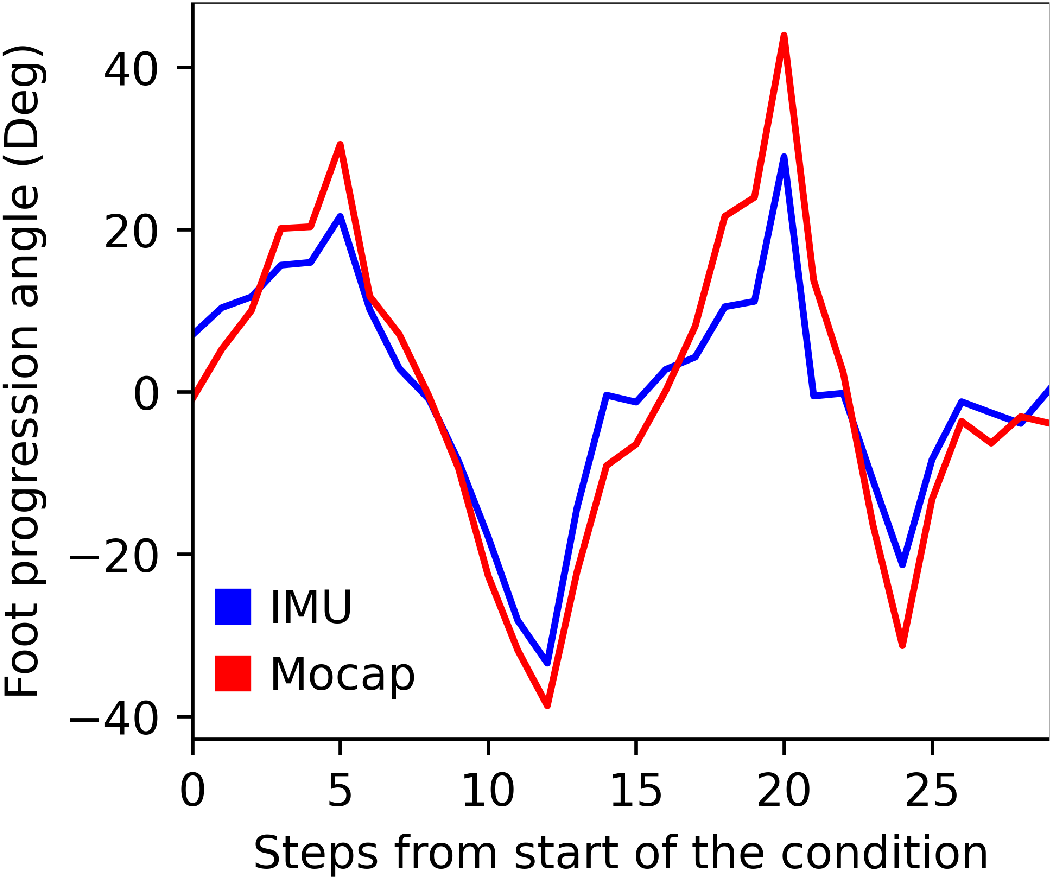
Comparison of foot progression angle computed by the IMU system and motion capture for one subject. The subject varied their foot progression angle with each step. The IMU system followed the same trends as the motion capture system, but with smaller values across most steps. The average RMSE across all steps was 7.6 degrees.

### B. Computational limitations

We evaluated the real-time computational limits of the OpenSenseRT system to understand the relationship between the number of sensors used to track body segments and the computation rate of the inverse kinematics solver. We evaluated the fastest computation rate for the system when using 1 to 14 sensors. We found the maximum computation rate when using a single core of the microcontroller had a line of best fit of 69 *—*4.8 *∗N*, with *N* denoting the number of IMU sensors used to compute the kinematics (Fig. 8A). This recommended maximum computation rate should be a guideline when customizing the number of IMU sensors for new applications. When computing kinematics using a single core, the microcontroller had a steady-state CPU temperature safely below the soft and hard temperature limits set by the operating system (Fig. 8B). Thus, the device can safely operate in this mode indefinitely without encountering computation limits due to heat. These computation experiments were performed indoors at a room temperature of approximately 22 degrees Celsius. We anticipate the single core mode of operation results reported here could be replicated in ambient temperatures up to 37 degrees Celsius without performance degradation.

**Fig. 8.**
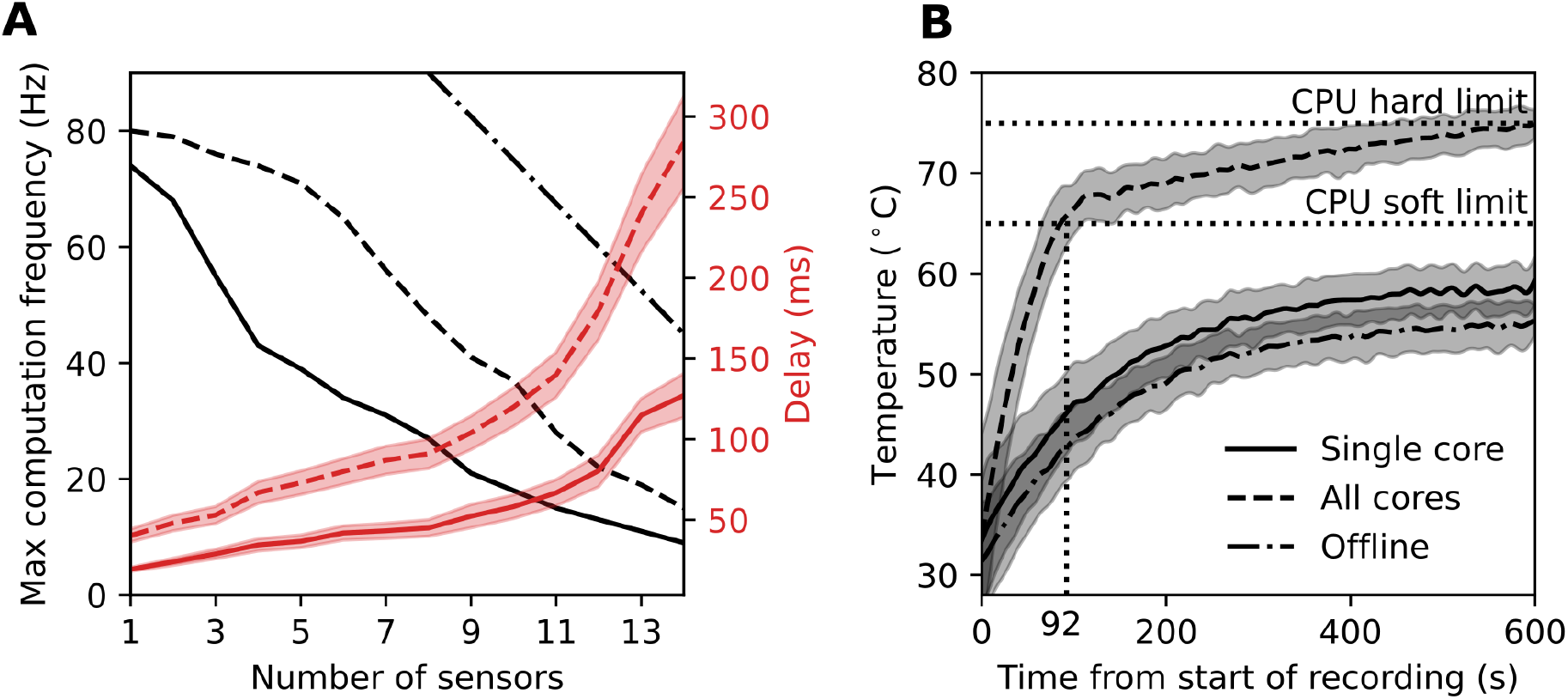
Computation and temperature limits of the microcontroller. (A) The maximum sampling rate and associated average delay depends on the number of sensors attached to the system and the mode of computation. We evaluate three modes of computation on the microcontroller using: a single core in real-time (solid line), all four cores in real-time (dashed line), and offline computation using a single core (dash-dot line). The sampling rates (black) and delays (red) were averaged over a one-minute recording. The error band represents one standard deviation across 5 trials. (B) The internal CPU processor temperature as a function of time was averaged for each of the 14 sensor configurations. The single core (solid line) and offline (dash-dot line) tests resulted in a safe steady-state temperature below any CPU limits.

A second real-time mode of operation utilized all cores on the microcontroller to increase the maximum computation at the expense of increased CPU temperature. The maximum computation rate in this mode had a line of best fit of 93 *—*5.6 *∗N*, where *N* denotes the number of IMU sensors. The CPU temperature reached the soft limit after 92 seconds and the hard limit after 10 minutes. The soft computation limit reduced the microcontroller performance and computation rate, preventing computation of joint kinematics in real time. Thus, using all cores may provide a faster real-time computation rate than a single core, but cannot be sustained without overheating the microcontroller.

An offline mode was tested by taking measurements from the IMUs in real-time and then computing the kinematics once the recording was finished. The computation in real-time was only limited by how fast the sensors could be sampled, allowing much faster offline computation rates of 150 *—* 7.5 *∗N*, where *N* is the number of IMU sensors. This mode offered sampling frequencies similar to other commercial systems [7] capable of capturing dynamic activities such as running.

## IV. Discussion

The OpenSenseRT system is the first open-source, low-cost system to estimate kinematics for a range of different movements, in real time. We computed and analyzed kinematics for 30 different degrees of freedom, including upper-and lower-extremity joint kinematics, with an average RMSE of 5 degrees compared to optical motion capture. The OpenSenseRT system provides flexibility, with high-frequency sampling, real-time or offline kinematics computation, and access to the modeling and analysis tools available in OpenSim, which we demonstrate by computing foot progression angle. The system provides technology to estimate joint kinematics in free-living conditions, which will help translate clinical tools into the real-world and enable large-scale and long-term biomechanics studies.

### A. Evaluating the accuracy of IMU motion capture

Joint kinematics estimated by the OpenSenseRT system had an average error of 5 degrees RMSE across lower and upper-extremity joints compared to optical motion capture, an acceptable error for use in many clinical and research applications [28]. Accuracy requirements will vary and should be validated for a specific clinical application before using the OpenSenseRT system to evaluate clinical outcomes. The error bands of hip and knee flexion during walking had similar magnitudes for both the OpenSenseRT system and optical motion capture, indicating that estimated kinematics were similarly consistent for both approaches. We expect there to be differences between kinematics estimates with OpenSenseRT and optical motion capture because optical motion capture is known to have measurement errors, for example due to skin motion artifacts. The accuracy of the OpenSenseRT system in measuring lower-limb joint kinematics is within the range of a state-of-the-art commercial Xsens system [11]. The upper-limb accuracy is within the range of other IMU systems that compute kinematics offline [29], [30].

### B. Feasibility for estimating kinematics during real-world use

The OpenSenseRT system meets usability and technical specifications to effectively estimate kinematics in real-world settings. The system is designed for continual monitoring applications: it is light weight to minimize fatigue, it has a rechargeable battery capable of recording for most of a day, it can store days or weeks of kinematic estimates in memory, and it allows for recordings to be manually started and stopped with the press of a button.The system is also meant to be easily reproducible–the device consists of low-cost and off-the-shelf components that can be assembled in approximately 20 minutes using common tools without soldering or coding [18].

The error in joint kinematic estimates drifted over time as expected, since our sensor fusion approach did not use magnetometer information. When using the OpenSenseRT system for extended recordings, we suggest recalibrating the system every 3 minutes to keep the average error below the recommended 5 degrees. For example, in an all day monitoring experiment the OpenSenseRT system could be automatically recalibrated whenever quiet standing is detected. Standing detection could be based on thresholding raw IMU accelerometer measurements to ensure the limbs are oriented in the same direction and not moving.

### C. Extending the system to custom applications

The OpenSenseRT relies on open-source software and hardware for transparency and extension to new applications. The software builds upon OpenSim [19], a software framework with well documented musculoskeletal analysis and simulation tools that can be integrated for future uses, such as computing muscle-tendon lengths and velocities. Although the use of a physiological skeletal model can be more computationally intensive than other approaches for computing kinematics, it allows kinematics estimates to be constrained to realistic joint limits and provides access to other biomechanics tools that rely on musculoskeletal modeling [19]. The open-source software also allows for custom additions to compute spatiotemporal metrics such as gait cycle duration (e.g., using the approach of [31]), which are important for clinical analysis of lower-limb motion. The OpenSenseRT provides hardware flexibility by allowing additional sensors, such as pressure insoles or electromyography sensors, to be connected to the microcontroller and open-source hardware, enabling customized measurements.

### D. Limitations

We have benchmarked the OpenSenseRT system’s basic functionality, but additional work is required to validate the system for more challenging applications, such as activities that require measuring both the upper- and lower-limbs or dynamic activities with a large range of motion or high speeds, such as squatting or sprinting. The computation rate is limited by the performance of the microcontroller, which prevents fast computation rates with many sensors, making a task like estimating full-body kinematics during running challenging. The CPU speed may be increased by overclocking the microcontroller or adding additional computation sources, such as a USB accelerator.

Using all cores of the microcontroller significantly increased the computation rate, but would require additional cooling to prevent overheating.

The accuracy of the OpenSenseRT system is dependent on the initial calibration which requires firmly and precisely attaching the IMUs to each body segment, which could pose challenges for real-world use. Methods for simplifying the calibration process and generalizing to any initial pose would allow for easier recalibration. It may also be possible to automate the calibration process when common poses such as standing occur naturally, streamlining long recordings. In order to accurately capture motion, the IMUs need to be firmly attached to the body as any shift in their placement accumulates error in the joint kinematics over time. Integrating the IMUs into clothes may also improve consistency for monitoring applications over extended periods of time.

### E. Conclusion

OpenSenseRT is an open-source, low-cost, easy-to-assemble, and customizable system for computing real-time kinematics. This system will enable researchers to translate biomechanics, rehabilitation, and sports performance research that rely on kinematics into real-world solutions. The system’s accuracy meets clinical standards for several common use cases in biomechanics and rehabilitation research. We have also documented and openly shared all hardware and software to allow other researchers to use the system in a wide range of new applications that leverage the biomechanics software tools included in OpenSim. For example, wearable and real-time joint kinematics will allow research to test biofeedback for gait retraining or sports performance outside of the lab, and the low cost could enable large-scale and longitudinal collection of free-living biomechanics data. The accessibility of this technology could also reduce the financial and technological barriers to perform state-of-the-art quantitative biomechanical analysis in low-income or developing countries with limited access to laboratory equipment, offering tools to improve musculoskeletal health on a global scale.

## Supporting information

Video demonstration

## ACKNOWLEDGMENT

The authors would like to acknowledge the staff of the Human Performance Lab for supporting the equipment used for the experimental validation of this work.

